# Nutrient deprivation differentially affects gene expression, immunity, and pathogen susceptibility across symbiotic states in a model cnidarian

**DOI:** 10.1101/2023.07.30.551141

**Authors:** Maria Valadez-Ingersoll, Pablo J. Aguirre Carrión, Caoimhe A. Bodnar, Niharika A. Desai, Thomas D. Gilmore, Sarah W. Davies

## Abstract

Mutualistic symbioses between cnidarians and photosynthetic algae are modulated by complex interactions between host immunity and environmental conditions. Here, we investigate how symbiosis interacts with nutrient limitation to influence gene expression and stress response programming in the sea anemone *Exaiptasia pallida* (Aiptasia). Transcriptomic responses to starvation were similar between symbiotic and aposymbiotic Aiptasia; however, aposymbiotic anemone responses were stronger. Starved Aiptasia of both symbiotic states exhibited increased protein levels of immune-related transcription factor NF-κB, its associated gene pathways, and putative target genes. However, this starvation-induced increase in NF-κB only correlated with increased immunity in symbiotic anemones. Furthermore, starvation had opposite effects on Aiptasia susceptibility to pathogen and oxidative stress challenges, suggesting distinct energetic priorities under nutrient scarce conditions. Finally, when we compared starvation responses in Aiptasia to those of a facultative coral and nonsymbiotic anemone, “defense” responses were similarly regulated in Aiptasia and the facultative coral, but not in the nonsymbiotic anemone. This pattern suggests that capacity for symbiosis influences immune responses in cnidarians. In summary, expression of certain immune pathways – including NF-κB – does not necessarily predict susceptibility to pathogens, highlighting the complexities of cnidarian immunity and the influence of symbiosis under varying energetic demands.

## Introduction

Reef-building corals are assemblages of cnidarian host cells, intracellular photosynthetic algae (Family Symbiodiniaceae), and a complex microbiome^1, 2^. These symbiotic relationships enable corals to flourish in nutrient-poor waters^3^. In a healthy cnidarian-algal symbiosis, the symbiont provides fixed carbon photosynthates to the host while the host provides CO_2_ and other essential nutrients (*e.g.,* nitrogen and phosphorus) to the symbiont^4–6^. Corals and the complex ecosystems they engineer are globally threatened by anthropogenic climate change and associated environmental perturbations including rising sea surface temperatures, increased exposure to disease, ocean acidification, oxygen limitation, and changes in nutrient availability^7, 8^. Understanding the mechanisms underlying this symbiosis is critical for predicting how cnidarians will be affected by accelerating anthropogenic change.

Previous work has shown that the negative impacts of climate change on corals can be mitigated by nutrient acquisition from the environment through heterotrophy^9–12^. However, the reliance of the host on either form of nutrient acquisition – prey capture (heterotrophy) or translocation from the algal symbiont – is variable across taxa and environments^13, 14^. On one extreme, cnidarians living in tropical, low-nutrient environments form obligate associations with their symbiotic algae. If this symbiosis is lost (termed “bleaching”), the host will eventually starve if it cannot buffer this energetic loss with heterotrophy^8^. In bleached corals, carbon incorporation from increased rates of heterotrophy is an important mechanism for recovery^15, 16^. Facultatively symbiotic cnidarians also exist; these species tend to live in higher nutrient environments and can acquire nutrients from a combination of their endosymbiotic algae and heterotrophy. This nutrient flexibility facilitates a range of states from symbiotic to aposymbiotic (extremely low levels of symbionts) depending on environmental conditions^17–19^. Finally, some cnidarians are nonsymbiotic, *i.e.,* they never form symbiotic relationships with photosynthetic algae and rely solely on heterotrophy^18^. This continuum of symbiotic strategies might ultimately influence the how cnidarians respond to environmental challenges.

Energetic priorities are likely to differ across symbiotic states and may further shift under variable nutrient conditions. Previous research has demonstrated that the nonsymbiotic sea anemone *Nematostella vectensis* is more susceptible to the effects of bacterial infection under nutrient limitation, which was associated with reduced expression of known innate immune pathways^20^. In obligate and facultative symbioses, maintaining the algal symbiotic relationship, even under ambient conditions, requires tradeoffs for hosts. Much of the research on these tradeoffs has been done in facultatively symbiotic cnidarian models, such as the sea anemone *Exaiptasia pallida* (Aiptasia)^21–23^. First, hosts must have the capacity to regulate symbiont cell density to control overcrowding, which becomes especially important when nutrient availability changes^24^. Previous work has shown that when symbiotic Aiptasia are starved, symbiont densities decrease by up to 50%, likely due to inter-algal competition for decreased host-derived nutrients^25^. Second, symbiotic hosts must balance allowing foreign algal cells to live intracellularly while also maintaining defense against harmful pathogens^26^. In general, research in symbiotic cnidarians has demonstrated that the establishment and maintenance of symbiosis requires downregulation of certain innate immune system pathways, including the transcription factor NF-κB pathway^27–29^. Thus, in symbiotic cnidarians, the nutritional benefits conferred by symbiosis are at a potential cost to the host’s defense against pathogens.

Field research has shown that symbiont loss during episodes of coral bleaching can confer heightened, though transient, protection against harmful pathogens^30^. However, stress responses are complex, and symbiotic cnidarians have also been shown to have higher levels of antioxidant enzymes (*i.e.,* catalase and superoxide dismutase) than aposymbiotic cnidarians, suggesting greater protection against oxidative stress in symbiotic animals^31–34^. These results suggest that NF-κB and oxidative stress response pathways are regulated differently in cnidarians. Taken together, two central themes of cnidarian symbiosis have emerged: 1) symbiosis modulates host immunity, and 2) heterotrophy modulates symbiosis. However, the influence of symbiosis and nutrition on immunity, and the distinct regulatory pathways of the broader immune system, remain largely unexplored, but may provide insight for understanding cnidarian stress responses.

Here, we have characterized the effects of nutrient limitation on gene expression in the facultatively symbiotic sea anemone Aiptasia when it is associated with algal symbionts (symbiotic/sym) or not (aposymbiotic/apo). We used gene expression data to investigate how starvation-induced modulation of immune and oxidative stress pathways impacts cnidarian susceptibility to pathogen and oxidative stress challenges. Finally, we compared starvation-induced gene expression patterns in Aiptasia to previously obtained gene expression patterns under starvation in a facultatively symbiotic stony coral (*Oculina arbuscula*) and a nonsymbiotic sea anemone (*Nematostella vectensis*). This comparative approach showed the evolutionary underpinnings of molecular responses to starvation in cnidarians and how symbiosis with Symbiodiniaceae algae can influence these responses. Our results suggest that, in facultatively symbiotic cnidarians, there are distinct molecular regulatory mechanisms for oxidative stress, the pathogen-responsive innate immune response, and the symbiont-responsive innate immune response. Furthermore, under nutrient limitation, energetic priorities within the immune system differ between symbiotic states and by the ability of a cnidarian species to form symbiosis with Symbiodiniaceae.

## Materials and Methods

### Aiptasia husbandry and experimental design

Symbiotic (partnered with *Symbiodinium linucheae*) and aposymbiotic adult *Exaiptasia pallida* (Aiptasia) anemones of clonal strain CC7^35^ were maintained in 35 ppt artificial seawater (ASW, Instant Ocean) in polycarbonate tanks at 25 ℃ under a 12 h:12 h light:dark cycle (white fluorescent light at 25 µmol photons/m^2^/s). Aposymbiotic Aiptasia were generated via menthol bleaching at least three months prior to experimentation as described previously^36^. Aposymbiotic Aiptasia were maintained in darkness and aposymbiotic status was confirmed by lack of symbiont autofluorescence under fluorescence microscopy (Leica M165 FC). Anemones were fed three times per week with freshly hatched *Artemia* nauplii, and water changes were performed twice weekly.

To generate clonal pairs, single anemones were placed into individual wells of a 6-well plate in 10 ml ASW. Anemones were relaxed on ice for 10 min, then bisected along the oral-aboral axis. After at least two weeks, bisected anemones were determined to be healed by visual confirmation of complete regeneration of the oral/tentacle disc. During recovery, anemones were maintained at 25 ℃ with symbiotic pairs under a 12 h:12 h light:dark cycle and aposymbiotic pairs under darkness. All clonal pairs were fed freshly hatched *Artemia* nauplii, and water changes were performed three times per week during regeneration.

Following regeneration, each anemone was placed into an independent well with 10 ml of 0.2 μM-filter sterilized ASW (FSW). Three days prior to use, aposymbiotic clonal pairs were moved to a 12 h:12 h light:dark cycle for light acclimation. During the course of the two-week treatment period, anemones were maintained under a 12 h:12 h light:dark cycle with 25 µmol photons/m^2^/s. For starvation experiments, one individual of each clonal pair was given 30 μl of *Artemia* nauplii three times per week (“fed” control), while the other was not fed (“starved”). To mimic feeding in starved anemones, 30 μl of FSW was pipetted onto all “starved” individuals. Water changes were performed twice weekly with 10 ml of FSW. After two weeks, anemones were preserved or processed for downstream analyses. Clonal pairs were not used for susceptibility trials, but the feeding/starvation regimen with non-bisected aposymbiotic and symbiotic CC7 Aiptasia used was identical to methods described above. The experimental design is shown in **Figure 1**.

**Figure 1.**
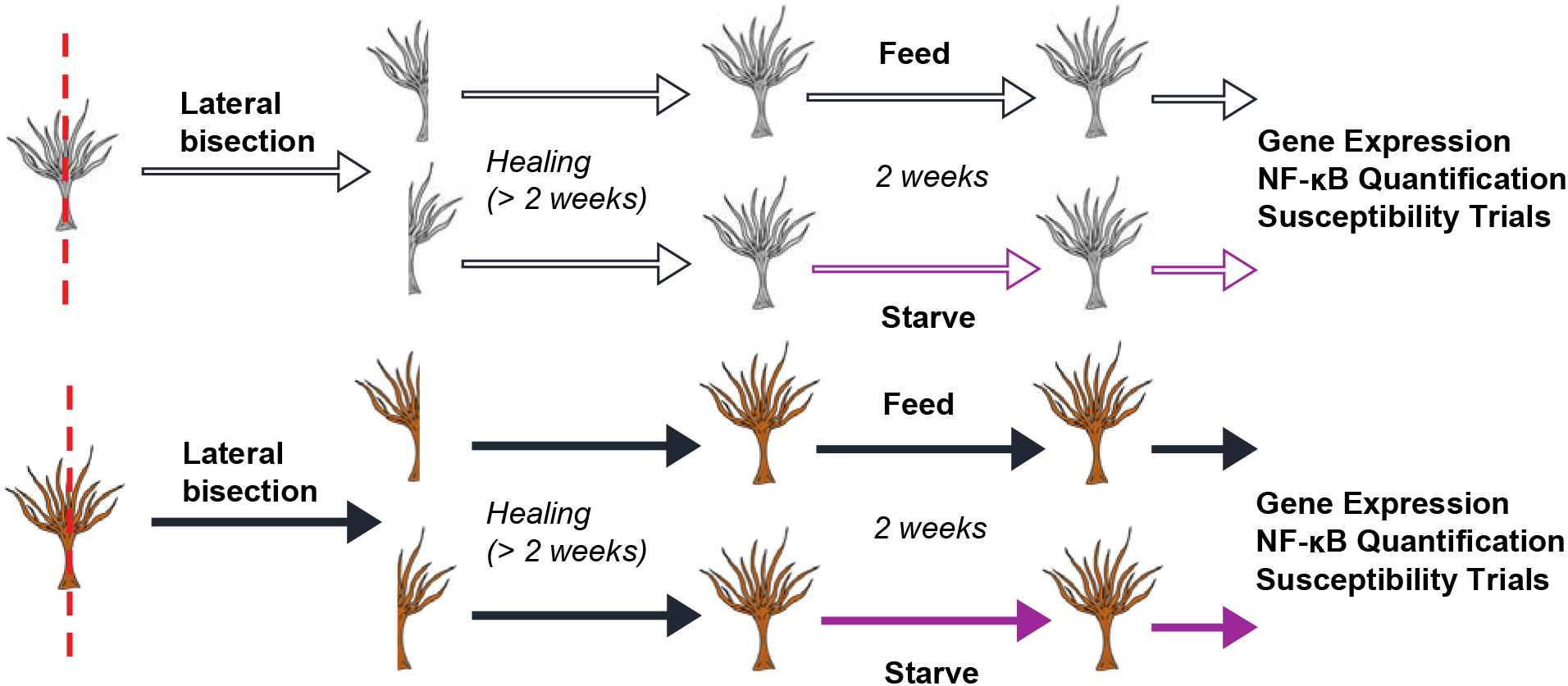
Experimental design. Aposymbiotic (top, grey) and symbiotic (bottom, brown) anemones were bisected to create clonal pairs. Halves were separated and allowed to heal for at least two weeks. After healing, one individual of each clonal pair was fed regularly and the other was starved for two weeks. Tissue of each individual was then preserved for subsequent analysis. Anemones destined for susceptibility trials were not cloned, but the same feeding/starvation conditions were employed.

### RNA extraction and TagSeq processing

After two weeks of treatment, five fed/starved clonal pairs of symbiotic and aposymbiotic Aiptasia (n=20) were flash frozen and preserved in 100% ethanol. Total RNA was extracted from each anemone by grinding whole anemones with plastic pestles during tissue lysis and centrifuging at 13,000 rpm for 15 min at 4 ℃. Supernatant was processed by the RNAqueous™ Total RNA Isolation Kit (Invitrogen) per manufacturer’s instructions. RNA quantity and integrity were assessed using a NanoDrop ND-1000 Spectrophotometer. Samples were normalized to 11.4 ng/µl in 30 µl and submitted to the University of Texas at Austin Genomic Sequencing and Analysis Facility for TagSeq library preparation and sequencing on a NovaSeq6000 SR100.

Raw fastq reads were processed following an established bioinformatics pipeline (https://github.com/z0on/tag-based_RNAseq). Briefly, fastx_toolkit was used to trim adapters and poly(A)^+^ tails, PCR duplicates were removed, and short (<20 bp) and low-quality reads (quality score of ≤20) were discarded^37^. Cleaned reads were mapped to concatenated Aiptasia^38^ and *Symbiodinium linucheae*^39^ transcriptomes using Bowtie2^40^. A raw counts file containing all genes in each sample mapping exclusively to the Aiptasia reference was used in downstream analyses (**Supplementary Dataset 1**). Symbiont-specific genes mapping to the *S. linucheae* reference were not analyzed due to low counts.

### Analysis of gene expression under starvation in Aiptasia

All analyses were performed in R v.4.3.0^41^. DESeq2 (v.1.36.0)^4^ was used to identify differentially expressed genes (DEGs) between fed and starved treatments in both symbiotic and aposymbiotic anemones. First, DESeq2 modeled all samples together by treatment (ApoFed, ApoStarved, SymFed, SymStarved). ArrayQualityMetrics (v.3.52.0)^43^ tested for outliers. No outliers were detected. Next, individual contrasts between fed and starved anemones within each symbiotic state were performed to identify DEGs (FDR-adjusted-p value of <0.05). Data were rlog-normalized and overall expression was compared through Principal Component Analysis (PCA) implemented in vegan (v.2.6-2)^44^. Gene expression similarity was tested across samples using PERMANOVA implemented with the pairwiseAdonis package v.0.4^45^. Next, gene expression plasticity of symbiotic and aposymbiotic Aiptasia under starvation was estimated as a function of the distance of each point in the treatment group (starved) relative to the location of the centroid in the control group (fed) in PCA space using the first two principal components^46, 47^. Gene expression plasticity for symbiotic and aposymbiotic Aiptasia was then compared using an analysis of variance. VennDiagram (v.1.7.3)^48^ compared total DEGs under starvation (FDR-p-values of <0.05) in symbiotic and aposymbiotic anemones to identify DEGs shared between symbiotic states under starvation, as well as DEGs unique to the starvation responses of each symbiotic state. Expression levels of shared DEGs were correlated using a linear model of log-fold change between starved vs fed anemones across symbiotic and aposymbiotic states.

Functional gene ontology (GO) enrichment analyses of shared DEGs and of DEGs unique to each symbiotic state under starvation were performed using Fisher Exact Tests (DEG presence/absence)^49^ to identify enrichment differences across these unique sets of genes in the ‘Biological Process’ (BP) Gene Ontology (GO) division^49^. Next, enrichment of Aiptasia gene pathways under starvation in symbiotic and aposymbiotic Aiptasia was independently investigated by GO enrichment analysis using the Mann-Whitney U tests (GO-MWU)^49^ on all genes. Significantly enriched GO terms (FDR-p-value of <0.1 from GO MWU tests) for both types of GO analyses (Fisher’s Exact and Mann-Whitney U tests) were plotted in dendrograms. We then identified significantly over and underrepresented BP GO terms under starvation that were explicitly involved in the NF-κB pathway (**Table 1**) and superoxide or reactive oxidative stress (ROS) response pathways (**Table 2**). A complete list of significantly enriched GO terms for all analyses are provided in **Supplementary Datasets 2 and 3**. Significant DEGs (FDR-p-value of <0.05) belonging to NF-κB and ROS GO terms of interest were visualized for symbiotic and aposymbiotic samples using pheatmap v.1.0.12^50^. Finally, we used a published dataset of heat-responsive Aiptasia genes (significantly upregulated after three hours of heat shock at 34℃) with three or more NF-κB binding sites in the promoter region (*i.e.*, 500 bp upstream of transcription start site)^51^ to compare to our data to identify possible NF-κB target genes that were significantly upregulated (FDR-p-value of <0.05) following starvation in symbiotic and aposymbiotic Aiptasia. Expression patterns were then visualized using pheatmap v.1.0.12^50^.

**Table 1.**
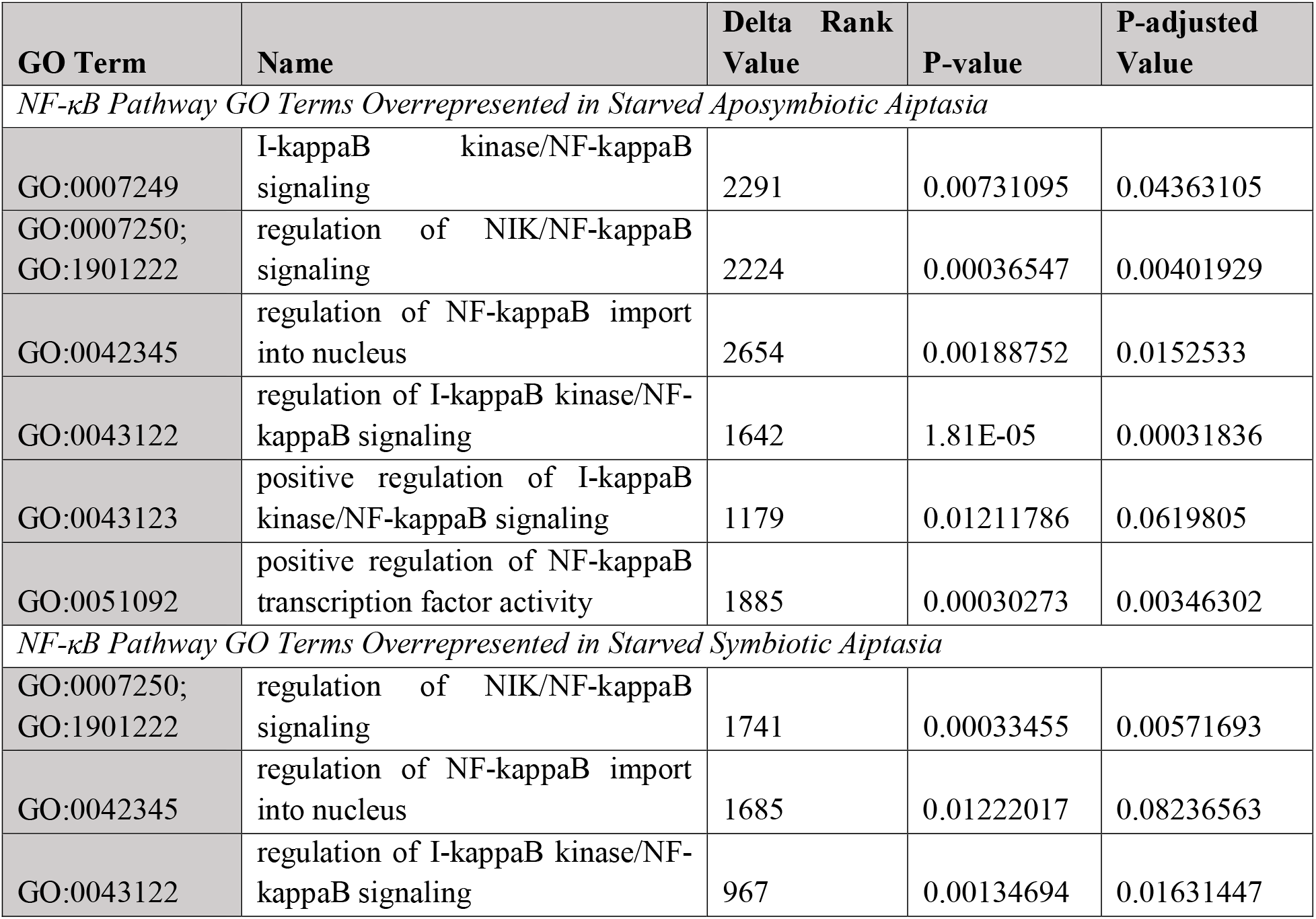
Biological Process Gene Ontology (GO) terms involved in the NF-κB pathway that are significantly overrepresented following starvation in symbiotic and aposymbiotic Aiptasia.

**Table 2.** Biological Process Gene Ontology (GO) terms involved in superoxide or reactive oxidative stress response pathways (ROS response) that are significantly underrepresented following starvation in symbiotic and aposymbiotic Aiptasia.

| GO Term | Name | Delta Rank Value | P-value | P-adjusted Value |
| --- | --- | --- | --- | --- |
| *ROS Response GO Terms Underrepresented in Starved Aposymbiotic Aiptasia* |  |  |  |  |
| GO:0000303;<br>GO:0019430;<br>GO:0071451 | response to superoxide | -3828 | 0.00077273 | 0.00745429 |
| GO:0072593 | reactive oxygen species metabolic process | -1186 | 0.01328871 | 0.06626742 |
| *ROS Response GO Terms Underrepresented in Starved Symbiotic Aiptasia* |  |  |  |  |
| GO:0000303;<br>GO:0019430;<br>GO:0071451 | response to superoxide | -2172 | 0.00718584 | 0.055403 |
| GO:0006801 | superoxide metabolic process | -1512 | 0.01265835 | 0.08402683 |
| GO:0072593 | reactive oxygen species metabolic process | -1309 | 0.0010438 | 0.01347666 |

### Western blotting of NF-κB

Individual anemones (n=12) were homogenized in 60 µl of 2X SDS-sample buffer (0.125 M Tris-HCl [pH 6.8], 4.6% w/v SDS, 20% w/v glycerol, 10% v/v β-mercaptoethanol, 0.2% w/v bromophenol blue) using a plastic pestle, followed by boiling for 10 min. Samples were centrifuged for 10 min at 13,000 rpm to remove cell debris, and the supernatant was collected. Protein isolation and Western blotting were performed as previously described^28, 52, 53^. Briefly, cell extracts were electrophoresed on a 7.5% SDS-polyacrylamide gel. Proteins were electrophoretically transferred to a nitrocellulose membrane in Transfer Buffer (20 mM Tris [pH 7.4], 150 mM glycine, 10% methanol). The membrane was blocked for 1 h at room temperature in TBST (10UmM Tris-HCl [pH 7.4], 150UmM NaCl, 0.05% v/v Tween 20 containing 5% non-fat powdered milk (Lab Scientific bioKEMIX, Inc.)). The membrane was then incubated overnight at 4 ℃ with primary rabbit-anti-Ap-NF-κB^28^ (diluted 1:5000 in TBST with 5% non-fat powdered milk). The membrane was washed five times with TBST and then incubated for 1 h with secondary goat-anti-rabbit-HRP conjugate (1:4000, Cell Signaling Technology) in TBST with 5% non-fat powdered milk. Membranes were washed four times with TBST, and then twice with TBS (10UmM Tris-HCl [pH 7.4], 150UmM NaCl). Immunoreactive proteins were detected with SuperSignal West Dura Extended Duration Substrate (Fisher Scientific) and imaged on a Sapphire Biomolecular Imager. Bands corresponding to Ap-NF-κB were normalized to total protein loaded in each lane using Ponceau S stain and ImageJ. For each clonal pair, total NF-κB protein in the starved individual was compared to that of its fed counterpart.

### Effect of starvation on pathogen- and H_2_O_2_-induced mortality in symbiotic and aposymbiotic Aiptasia

Pathogen susceptibility trials were performed as described previously^20^ with the general pathogen *Pseudomonas aeruginosa* strain Pa14 and the opportunistic coral pathogen *Serratia marcescens* (Smarc). Pa14 and Smarc susceptibility trials were each performed in duplicate under the following conditions: symbiotic fed (n=24), symbiotic starved (n=24), aposymbiotic fed (n=24), aposymbiotic starved (n=24), comprising a total of n=48 per condition and n=192 total anemones per pathogen trial. Briefly, anemones were fed or starved for two weeks, transferred to individual wells of a 24-well plate with 1 ml FSW, and acclimated at 27 °C for 24 h prior to infection. Single colonies of Pa14 and Smarc were cultured overnight at 37 ℃ in Luria Broth. Bacteria were washed three times with FSW and then diluted in FSW to an OD_600_ of approximately 0.1. FSW was then replaced with 1 ml of Pa14 (average of ∼ 3.94×10^8^ CFU/ml) or 1 ml of Smarc (average of ∼ 3.63×10^8^ CFU/ml). Infected anemones were maintained at 27 ℃ under a 12 h:12 h light:dark cycle and monitored for viability each day for four weeks. Aiptasia mortality was assessed by prevalent tissue lysis and by lack of response to stimulus (touch with sterile pipette tip and squirt with water). Symbiotic and aposymbiotic Aiptasia (n=20) were additionally incubated in FSW at 27 ℃ for the four-week monitoring period as experimental controls.

Susceptibility of fed versus starved Aiptasia to reactive oxygen species (ROS) was tested as previously described^54^. Briefly, ROS susceptibility trials were performed in duplicate (symbiotic fed (n=24), symbiotic starved (n=24), aposymbiotic fed (n=24), aposymbiotic starved (n=24), comprising a total of n=48 per condition and n=192 total anemones). Anemones were fed or starved for two weeks and then transferred to individual wells of a 24-well plate containing 1 ml of 0.008% (2.6 mM) H_2_O_2_ in FSW. Daily water changes with fresh 0.008% H_2_O_2_ in FSW were performed, and mortality was monitored daily for four weeks. Anemones were maintained at room temperature under a 12 h:12 h light:dark cycle.

### Survival analyses

For each condition, results from duplicate trials were combined and analysed together. The survival v.3.5-5 and survminer v.0.4.9 packages^55–57^ analyzed results of the pathogen and ROS susceptibility trials. For each stressor, Cox proportional hazards models were used to model the effect of treatment (symbiotic fed (SymFed), symbiotic starved (SymStarved), aposymbiotic fed (ApoFed), and aposymbiotic starved (ApoStarved)) on the survival probability. Following this model, we conducted four pairwise comparisons: SymFed to SymStarved, ApoFed to ApoStarved, SymFed to ApoFed, and SymStarved to ApoStarved.

### Meta-analysis of the starvation response across Cnidaria

A meta-analysis of gene expression responses to starvation across cnidarians was performed by comparing our data to previously published datasets generated from fed/starved *N. vectensis* (nonsymbiotic sea anemone)^20, 58^ and symbiotic *O. arbuscula* (facultatively symbiotic stony coral)^10^. This step first confirmed nutrient limitation under experimental starvation conditions across experiments and then contrasted differences in stress responses across cnidarian taxa. All over- or underrepresented BP GO terms under starvation (FDR-p-value of <0.1 from GO MWU) in two or more of the study species (*N. vectensis, O. arbuscula,* and Aiptasia) were identified. This approach revealed 101 GO terms from which 37 metabolism-related terms and 6 defense-related terms identified and these GO term delta rank values were compared across taxa and visualized in ggplot2 (v.3.4.0)^59^.

## Results

### Aposymbiotic Aiptasia exhibit stronger transcriptomic responses to starvation than symbiotic Aiptasia

To compare the effect of starvation on gene expression in symbiotic and aposymbiotic Aiptasia, the overall patterns of gene expression were compared across treatment groups (Symbiotic Fed, Symbiotic Starved, Aposymbiotic Fed, and Aposymbiotic Starved) through Principal Component Analysis (PCA). Based on PCA clustering, nutrition status was a stronger predictor of gene expression patterns (samples differentiated on PC1 explaining 30.8% of the variation) than symbiotic state (samples differentiated on PC2 explaining 11.9% of the variation); however, both factors were significant (p<0.001 for nutrition status and p=0.017 for symbiotic state, **Fig. 2A)**. The interaction between nutrition and symbiotic status was also significant (p=0.043, **Fig. 2A**). Aposymbiotic Aiptasia exhibited stronger transcriptomic responses to starvation than symbiotic Aiptasia, as revealed by higher gene expression plasticity under starvation (p<0.001, **Fig. 2B**). Furthermore, under starvation, aposymbiotic Aiptasia had a greater number of DEGs (FDR-p-value of <0.05) than symbiotic anemones: we identified 4,858 starvation-induced DEGs in aposymbiotic compared to 1,724 starvation-induced DEGs in symbiotic Aiptasia (**Fig. 2C**). A total of 1,460 starvation-induced DEGs were shared between symbiotic and aposymbiotic Aiptasia, of which only 10 were expressed in opposite directions (**Fig. S1** and **Table S1**), showing the similarity of the starvation response between symbiotic states. Lastly, from Fisher Exact Tests, Biological Processes (BP) GO term enrichment of shared DEGs and symbiotic state-specific DEGs revealed enrichment of pathways involved in growth, metabolism, and response to oxidative stress (**Supplementary Dataset 2**). This observed BP GO term enrichment was exaggerated in aposymbiotic-unique DEGs; however, no clear functional differences across BP GO terms were observed between these categories (**Supplementary Dataset 2**).

**Figure 2.**
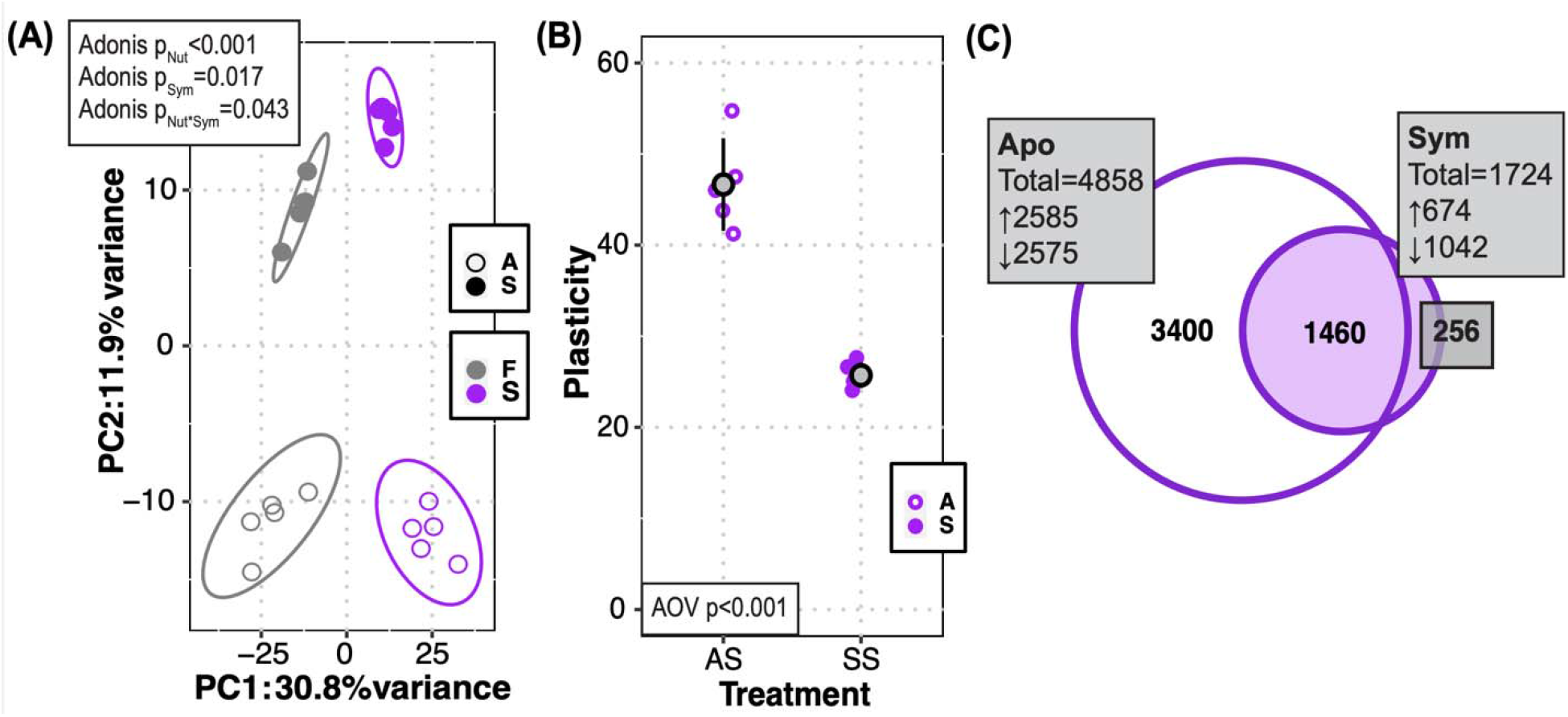
Starvation elicits shifts in whole-transcriptome gene expression and stronger transcriptomic responses in aposymbiotic compared to symbiotic Aiptasia. **(A)** Principal component analysis (PCA) of log2 transformed counts for aposymbiotic (A, open circles) and symbiotic (S, closed circles) Aiptasia after two weeks of feeding (F, grey) or starvation (S, purple). PERMANOVA results for the effect of nutrition status (p_Nut_), symbiotic status (p_Sym_), and the interaction between nutrition and symbiotic state (p_Nut*Sym_) are included. **(B)** Gene expression plasticity for aposymbiotic Aiptasia (A, purple open circles) and symbiotic Aiptasia (S, purple closed circles) following two weeks of starvation was calculated from the PCA using the distance of each starved sample relative to the location of the centroid in the fed group. Grey circles show the mean and black bars indicate standard deviation. Significance value by analysis of variance (AOV) is included. **(C)** Venn Diagram of all differentially expressed genes (DEGs) as identified via DESeq2 with an FDR adjusted-p-value of <0.05 for symbiotic (Sym) and aposymbiotic (Apo) Aiptasia after two weeks of starvation compared to two weeks of regular feeding. At an adjusted p-value of <0.05, there were a total of 4,858 DEGs in aposymbioti Aiptasia and a total of 1,724 DEGs in symbiotic Aiptasia. 1,460 DEGs were shared between Sym and Apo Aiptasia under starvation.

### Overrepresentation of immune pathways and underrepresentation of oxidative stress response pathways following starvation regardless of symbiotic state

To understand the functional drivers of the Aiptasia starvation response, we probed the over- and underrepresented GO terms detected through BP GO term enrichment analysis of both symbiotic and aposymbiotic Aiptasia under starvation (**Supplementary Dataset 3**). This analysis revealed strong signatures of enrichment of metabolism, immunity, and response to reactive oxygen species following starvation in both symbiotic and aposymbiotic states. GO terms associated with the innate immune system–-specifically the NF-κB pathway (*e.g.*, “regulation of I-kappaB kinase/NF-kappaB signaling” GO:0043122)–were overrepresented under starvation in both symbiotic and aposymbiotic conditions (**Table 1**). Additionally, general immune and stress response terms (referred to as “ROS response” hereafter; *e.g.*, “reactive oxygen species metabolic process” GO:0072593) were underrepresented under starvation in both symbiotic and aposymbiotic conditions (**Table 2**).

From the overrepresented NF-κB pathway GO terms, we identified genes that were significantly upregulated under starvation in both symbiotic states (FDR-p-value of <0.05), which included two Tumor Necrosis Factor Receptor-Associated Factor homologs (TRAF4 and TRAF6) and a B-cell Lymphoma 3 (BCL3) homolog (**Fig. 3A**). Among the NF-κB pathway genes that were significantly upregulated under starvation in aposymbiotic Aiptasia only, we noted an additional BCL3 homolog, multiple homologs of NF-κB response genes (*e.g.*, Tumor Necrosis Factor alpha-induced protein 3 (TNFAIP3)), and the NF-κB gene itself (**Fig. 3A**). However, a lesser number of NF-κB pathway genes were downregulated following starvation (*e.g.*, CANT1).

**Figure 3.**
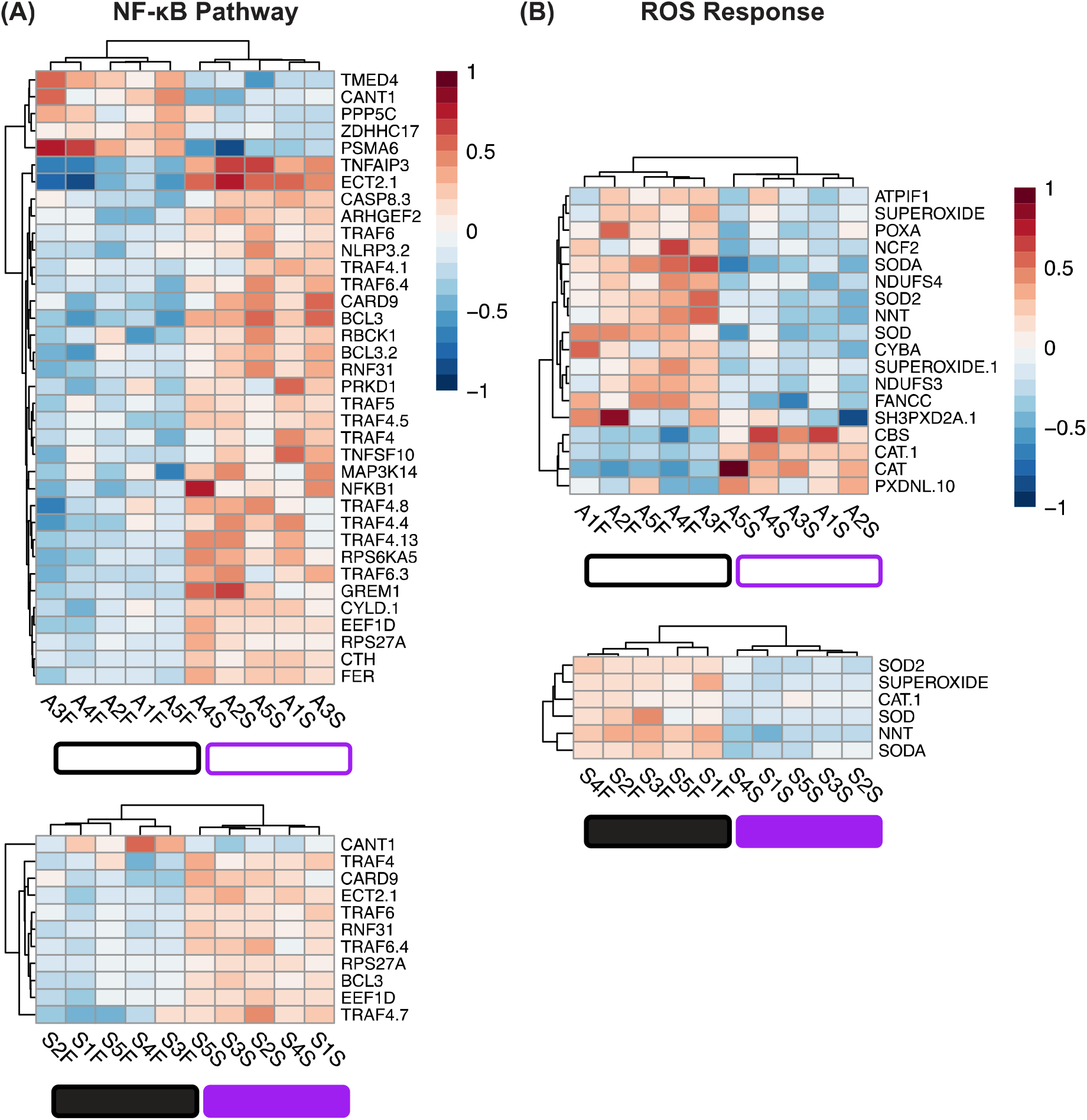
Differential regulation of NF-κB and ROS stress responses under starvation conditions in Aiptasia. **(A)** Significantly differentially expressed genes (FDR-adjusted-p value <0.05) in the NF-κB pathway (aposymbiotic clonal pairs top, symbiotic clonal pairs bottom). The NF-κB pathway was chosen to highlight a symbiosis-related immune system pathway. **(B)** Significantly differentially expressed genes (FDR-adjusted-p value <0.05) involved in the response to Reactive Oxygen Species (ROS; aposymbiotic clonal pairs top, symbiotic clonal pairs bottom). The ROS response was chosen to highlight a general cnidarian stress response. The colour scale of the heatmaps represents the log_2_ fold change of each gene (row) for each anemone (column) relative to the gene’s mean across all anemones where shades of red indicate upregulation and shades of blue indicate downregulation. In both panels, aposymbiotic anemones are represented by open boxes and symbiotic anemones are represented by closed boxes. Purple indicates starved samples and black indicates fed samples.

From the underrepresented GO terms associated with the ROS response, we identified genes that were significantly downregulated under starvation in both symbiotic and aposymbiotic Aiptasia (FDR-p-value of <0.05). These downregulated genes included four homologs for Superoxide Dismutase (SODA (a superoxide dismutase [Fe] homolog), SUPEROXIDE (a superoxide dismutase [Cu-Zn] homolog), SOD2 (a superoxide dismutase [Mn] homolog), and SOD (a putative superoxide dismutase [Cu-Zn] homolog)) (**Fig. 3B**). Among the oxidative stress response DEGs downregulated under starvation in aposymbiotic Aiptasia only, we identified an additional superoxide dismutase homolog (**Fig. 3B**). In aposymbiotic Aiptasia, some genes in the ROS Response pathway were upregulated following starvation (*e.g.*, the Catalase homolog CAT).

### Starvation leads to upregulation of NF-κB protein and putative NF-κB target genes in both symbiotic and aposymbiotic Aiptasia

Because NF-κB pathway GO terms and associated genes were enriched in starved Aiptasia, we sought to determine whether NF-κB protein levels were higher in starved anemones relative to fed ones. By Western blotting, we compared NF-κB protein levels in clonal pairs of fed and starved symbiotic and aposymbiotic Aiptasia. NF-κB protein levels were approximately 2.8 times higher in starved compared to fed symbiotic Aiptasia, and approximately 2.5 times higher in starved compared to fed aposymbiotic Aiptasia (**Fig. 4A**).

**Figure 4.**
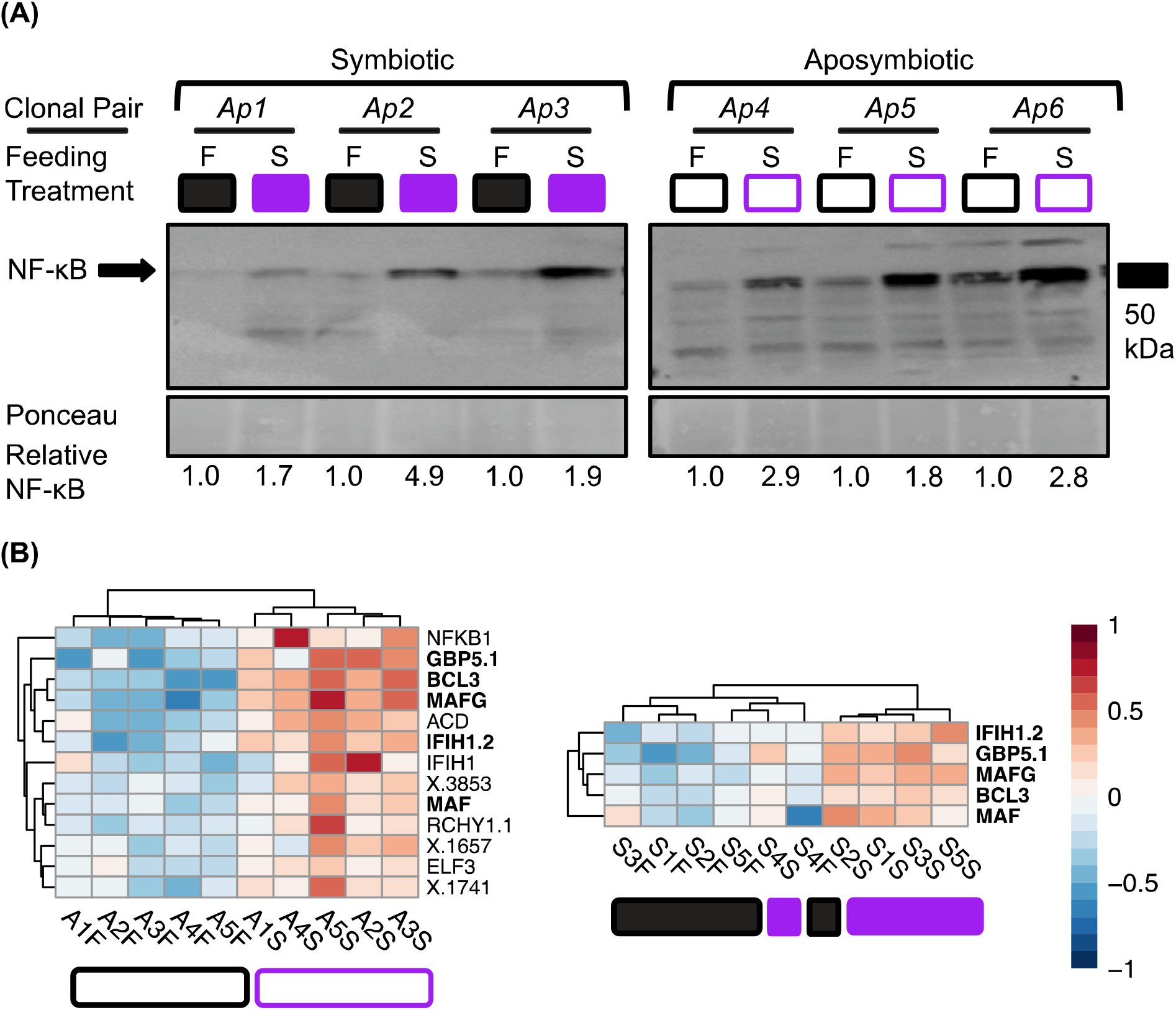
Upregulation of the NF-κB protein and putative NF-κB target genes under starvation in Aiptasia. **(A)** Western blot of NF-κB protein in symbiotic (left) and aposymbiotic (right) clonal pairs (Ap1-6) under fed (F) or starved (S) conditions. NF-κB protein levels were normalized to Ponceau staining and quantified with ImageJ. Values below the blots are normalized NF-κB protein levels relative to the fed Aiptasia of each clonal pair (1.0) (*e.g.,* F1/F1 and S1/F1 for Ap1). The location of the 50 kDa molecular weight marker is indicated to the right of the blots. **(B)** Significantly upregulated (FDR p-value <0.05) heat-responsive genes previously characterized as putative NF-κB target genes^51^ in aposymbiotic clonal pairs (left) and symbiotic clonal pairs (right). These genes have three or more NF-κB binding sites in the region 500 bp upstream of the transcription start site^51^ (Table S2 and Table S3). Bolded genes are shared between aposymbiotic and symbiotic states. The gene name “ACD” was shortened from “ACD_16C00100G0095”. Heatmap colour scale represents the log2 fold change of each gene (row) for each anemone (column) relative to the gene’s mean across all anemones with shades of red indicating upregulation and shades of blue indicating downregulation. In both panels, aposymbiotic anemones are represented by open boxes and symbiotic anemones are represented by closed boxes. Purple indicates starved samples and black indicates fed samples.

To further explore whether the NF-κB pathway was upregulated following starvation in symbiotic and aposymbiotic Aiptasia, we identified significantly upregulated genes under starvation (FDR-p-value of <0.05) that have been previously shown to be heat responsive and to contain at least three NF-κB binding sites in the region 500 bp upstream of the transcription start site (**Table S2 and Table S3**)^51^. In starved symbiotic and aposymbiotic Aiptasia, significantly upregulated putative NF-κB target genes included BCL3 and other genes implicated in innate immunity (*e.g.*, Interferon-induced helicase C domain-containing protein 1 (IFIH1) and Guanylate-binding protein 5 (GBP5); **Fig. 4B**). Starved aposymbiotic Aiptasia showed upregulation of several additional putative NF-κB target genes, including NF-κB itself and the ETS-related transcription factor ELF3 (**Fig. 4B**).

### Pathogen and ROS susceptibility is variable across different symbiotic and nutritional states

To determine whether the observed differential regulation of the immune and oxidative stress responses following starvation leads to altered susceptibility to immune and oxidative challenges, we performed pathogen and ROS exposure experiments. We found that anemones exposed to either four weeks of the bacterial pathogens Pa14 or Smarc showed differential mortality across symbiotic states (symbiotic or aposymbiotic) and nutritional status (fed or starved). Symbiotic Fed (SymFed) Aiptasia exposed to Pa14 had higher risk of mortality (lower survival) than Symbiotic Starved (SymStarved) Aiptasia (p<0.001) (**Fig. 5**). A similar result was also observed in SymFed and SymStarved Aiptasia exposed to Smarc (p<0.001) (**Fig. 5**). Additionally, Aposymbiotic Starved (ApoStarved) Aiptasia had a higher risk of mortality than SymStarved Aiptasia following exposure to both pathogens (p<0.001) (**Fig. 5**). However, in both pathogen challenges, starvation increased bacterial pathogenicity in aposymbiotic Aiptasia, as observed by the higher risk of mortality and lower survival probability in ApoStarved compared to Aposymbiotic Fed (ApoFed) Aiptasia (p<0.001 for Pa14, p<0.01 for Smarc) (**Fig. 5**). Finally, ApoFed had a consistently lower risk of mortality than SymFed following exposure to Pa14 (p<0.01) and Smarc (p<0.001) (**Fig. 5**). Overall, these results demonstrate that starvation is protective against pathogen-induced mortality in symbiotic, but not aposymbiotic, Aiptasia.

**Figure 5.**
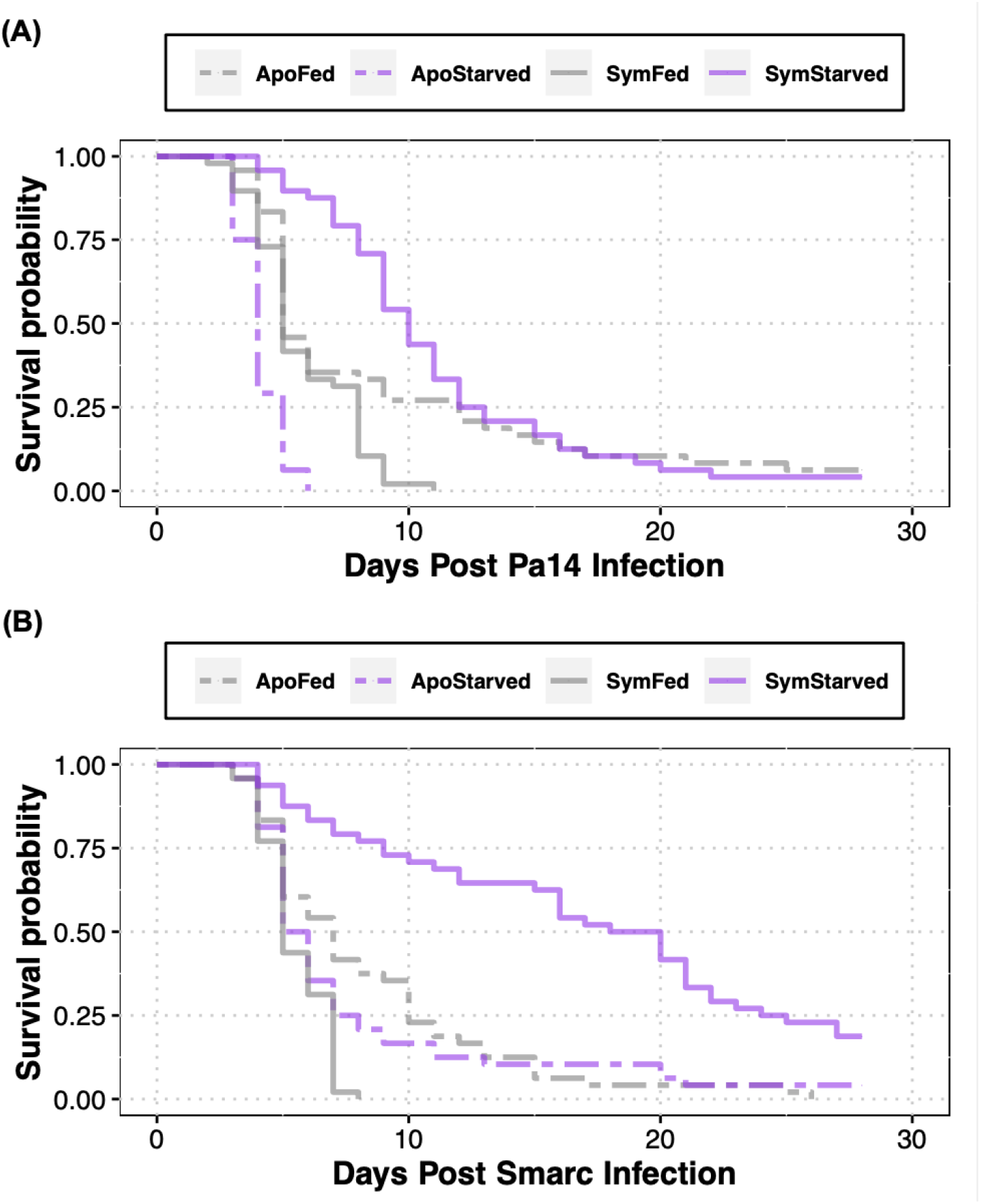
Starvation is protective against pathogen-induced mortality in symbiotic, but not aposymbiotic Aiptasia. Survival curves in fed and starved symbiotic and aposymbiotic Aiptasia exposed to 28 days of either *Pseudomonas aeruginosa* (Pa14) **(A)** or *Serratia marcescen* (Smarc) **(B)**.

When exposed to four weeks of 0.008% H_2_O_2_, both symbiotic and aposymbiotic anemones showed lower survival probabilities under starvation conditions as compared to fed Aiptasia exposed to H_2_O_2_ (p<0.001 for aposymbiotic and symbiotic) (**Fig. 6**).

**Figure 6.**
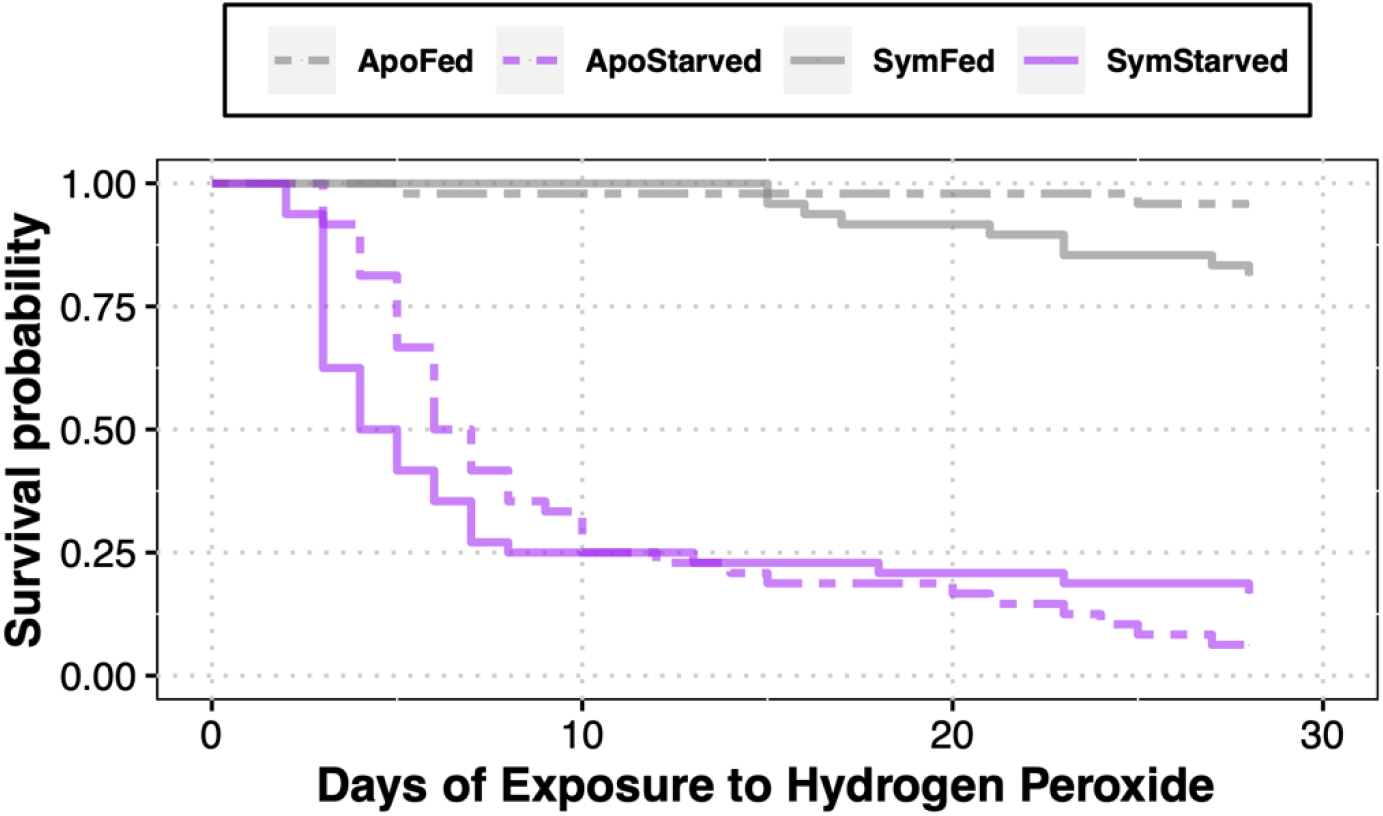
Starvation increases susceptibility to oxidative stress in symbiotic and aposymbiotic Aiptasia. Survival curves in fed and starved symbiotic and aposymbiotic Aiptasia exposed to 28 days of 0.008% H_2_O_2_.

### Among different cnidarians, starvation leads to similar changes in gene expression related to metabolism and oxidative stress, but not immune response

We performed a meta-analysis using previously published cnidarian starvation datasets to determine the degree of similarity in transcriptomic responses to starvation among different cnidarians. In symbiotic and aposymbiotic Aiptasia, nonsymbiotic *N. vectensis,* and symbiotic branches of the coral *O. arbuscula*, we observed downregulation of metabolic processes following starvation (**Fig. S2**). First, by comparing the delta rank values of all BP GO terms that were over- or underrepresented (FDR-p-value of <0.1) following starvation in at least two taxa, we observed extensive downregulation of terms associated with biosynthetic (*e.g.*, “carbohydrate biosynthetic process” GO:0016051), metabolic (*e.g*., “carbohydrate metabolic process” GO:0005975), and catabolic (*e.g.*, “carbohydrate catabolic process” GO:0016052) processes. These results indicate that starvation consistently restricts energetic supply to cause gene expression changes across different experiments, species, and symbiotic states (**Fig. 7A**).

**Figure 7.**
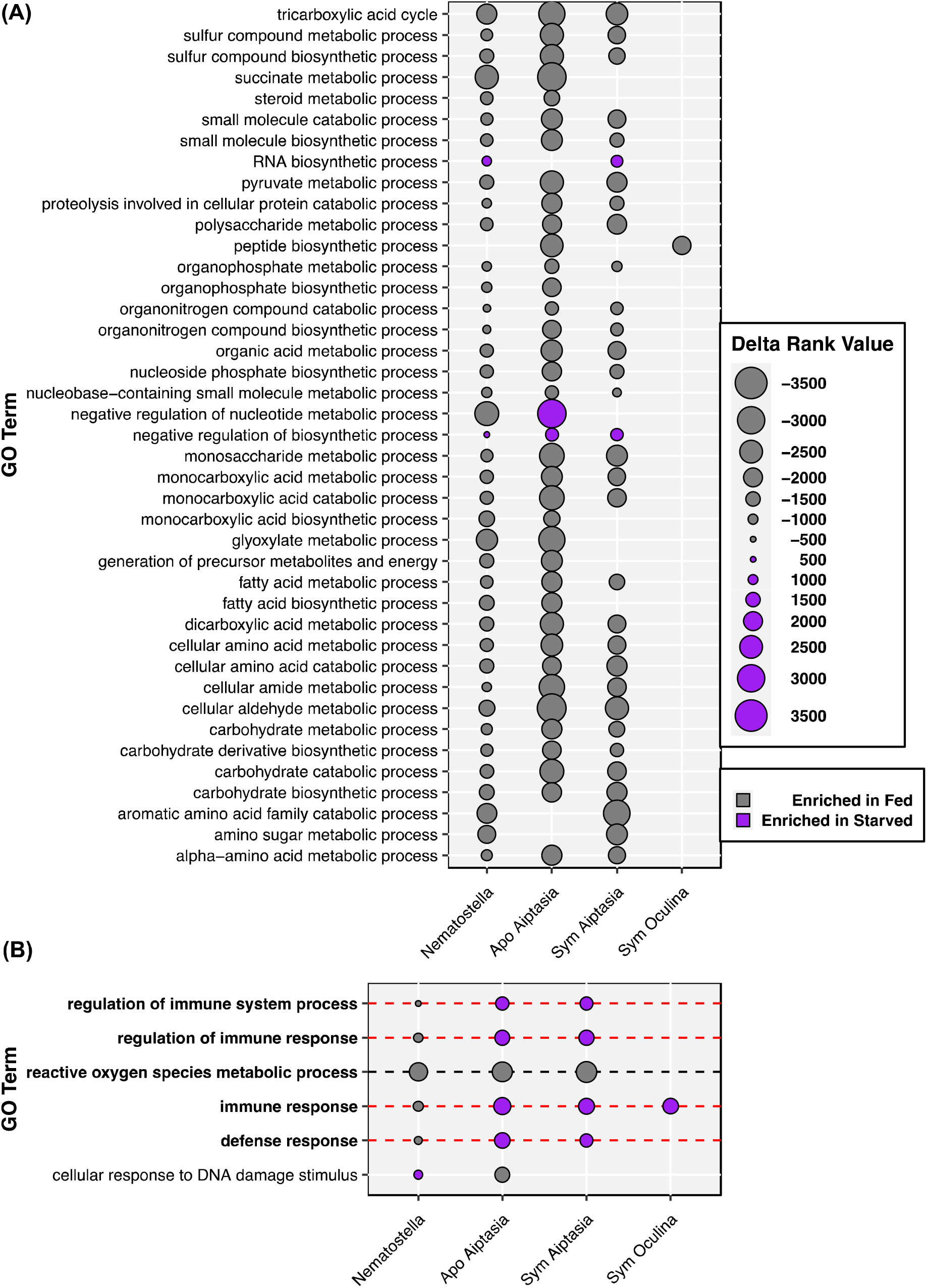
Gene Ontology (GO) term comparisons confirm nutrient limitation and reveal divergent regulation of immune and stress response pathways across Cnidaria. Delta rank values (represented by point size) were compared across starvation experiments in *Nematostella vectensis* (*Nematostella*^20^), aposymbiotic (Apo, this study) Aiptasia, symbiotic (Sym, this study) Aiptasia, and symbiotic *Oculina arbuscula* (Sym Oculina^10^). Terms were plotted if they were significantly enriched in fed (grey, negative delta rank values) or starved (purple, positive delta rank values) treatments in at least two comparisons. GO terms enriched in aposymbiotic and symbiotic Aiptasia, but not in either of the other species, were not included. (A) Shared GO terms that correspond to metabolism, catabolism, and biosynthesis were plotted to demonstrate their conserved downregulation under starvation across cnidarian taxa. (B) Shared GO terms that correspond to immunity, stress, and defense were plotted to demonstrate the differences in regulation of these pathways in *Nematostella* compared to Aiptasia and Oculina. Bold GO groups are those that are significantly enriched under starvation (either positively or negatively) in at least three systems. A red, dashed line indicates GO groups enriched in the opposite direction in *Nematostella* compared to at least two of the three Aiptasia and Oculina systems. A black, dashed line indicates GO groups enriched in the same direction in at least three systems.

GO enrichment analysis of starvation in symbiotic and aposymbiotic Aiptasia showed clear gene enrichment of immunity, stress, and defense (**Fig. S2**). To determine if these patterns were also observed in other cnidarians, we extracted enriched GO terms (FDR-p-value <0.1) associated with immunity, stress, and defense from the starvation experiments in *N. vectensis* and *O. arbuscula*. We found that many terms associated with broad immune system processes (*e.g.*, “regulation of immune system process” GO:0002684 and “defense response” GO:0006952) were enriched in the opposite direction in nonsymbiotic *N. vectensis* compared to species that are capable of symbiosis (Aiptasia and *O. arbuscula*). More specifically, the GO term “Immune Response” (GO:0006955) was overrepresented under starvation conditions in symbiotic Aiptasia, aposymbiotic Aiptasia, and symbiotic *O. arbuscula,* but was underrepresented in starved *N. vectensis*. However, GO terms associated with the oxidative stress response pathway (“reactive oxygen species metabolic process” GO:0072593) were underrepresented in starved *N. vectensis* along with starved symbiotic and aposymbiotic Aiptasia (these terms were not significant in *O. arbuscula*) (**Fig. 7B**). Overall, stress response pathways associated with ROS were downregulated under starvation across taxa, whereas cnidarians capable of symbiosis consistently upregulated immune system pathways under starvation.

## Discussion

Here, we describe the effects of nutrient limitation on gene expression, immunity, and stress responses in symbiotic and aposymbiotic anemones of the cnidarian model Aiptasia. We find that there is a core set of genes whose expression changes in response to starvation in both symbiotic and aposymbiotic anemones; however, there is also a large number of gene expression changes that occur only in aposymbiotic anemones. In both symbiotic and aposymbiotic anemones, starvation induces expression of the NF-κB transcription factor, as well as associated genes in the NF-κB pathway; however, starvation-induced upregulation of NF-κB is not necessarily predictive of organismal susceptibility to pathogen-induced mortality in Aiptasia. Furthermore, pathogen-specific immunity appears to be regulated independently from the Aiptasia oxidative stress response, suggesting that these two “defense” responses have distinct energetic priorities under nutrient scarce conditions. Finally, a meta-analysis comparing Aiptasia gene expression responses to starvation shows that cnidarian species – including Aiptasia, the non-symbiotic anemone *N. vectensis*, and the facultative coral *O. arbuscula* – vary in how they respond to starvation to a degree that suggests that starvation-induced gene expression and pathway responses are modulated by the animal’s ability to undergo symbiosis with Symbiodiniaceae algae.

One striking feature of our study was the strength of the transcriptomic response to starvation that occurred in aposymbiotic Aiptasia as compared to symbiotic Aiptasia. That is, starvation resulted in approximately three times as many DEGs in aposymbiotic anemones as compared to symbiotic anemones (4,858 vs 1,724), with aposymbiotic Aiptasia modifying the expression of approximately 18% of its transcriptome under starvation (4,858/∼27,000). Furthermore, although starvation was a stronger predictor of overall transcriptomic profiles than symbiotic state (supported by the 1,460 shared starvation-induced DEGs between symbiotic and aposymbiotic anemones), there were 3,400 starvation-induced DEGs that were unique to aposymbiotic Aiptasia, as compared to only 256 DEGs unique to symbiotic anemones. Therefore, only about 15% (256/1,724) of the DEGs in symbiotic anemones are unique, whereas ∼70% (3,400/4,858) are unique in aposymbiotic anemones. Pathway analysis of shared DEGs between aposymbiotic and symbiotic anemones under starvation revealed enrichment of processes involved in regulating growth, metabolism, and response to stress, a pattern that was amplified in aposymbiotic-specific DEGs. Although the effect of starvation on gene expression is stronger in aposymbiotic Aiptasia than symbiotic Aiptasia, these analyses reveal that the overall regulatory processes that are rewired under nutrient scarce conditions are similar between symbiotic states.

The stronger effects of starvation on aposymbiotic Aiptasia are likely because starved symbiotic Aiptasia can still obtain energy from their algal symbionts. Therefore, symbiotic Aiptasia may be buffered (by symbiont-derived energy sources) from the effects of starvation, whereas aposymbiotic Aiptasia must mount a broader gene expression response in order to survive starvation conditions. These results suggest that energetic reserves, whether from symbiotic or heterotrophic sources, provide a mechanism for protecting against the adverse impacts of environmental stress. Indeed, previous research has shown that corals that can compensate for the loss of symbiont-derived nutrients with heterotrophy are more resilient to heat-induced algal loss^11, 15, 16^. As such, in the absence of energetic reserves from either symbiotic or heterotrophic sources, cnidarians must extensively restructure their gene expression profiles to optimize energy use and prioritize gene pathways that promote survival.

We also found that starvation led to increased levels of NF-κB protein, as well as enrichment of NF-κB-related GO terms, associated pathway genes (*e.g.,* TRAF4, TRAF6, and TNFAIP3), and possible target genes (*e.g.,* BCL3, and GBP5), in both symbiotic and aposymbiotic Aiptasia. These results contrast with recent work in the nonsymbiotic anemone *N. vectensis*, where starvation led to reduced NF-κB protein and associated gene expression pathways^20^. Moreover, in *N. vectensis*, decreased NF-κB protein is correlated with increased susceptibility to bacterial pathogenesis^20, 60^. Although starvation-induced increases in NF-κB were correlated with decreased susceptibility to bacterial pathogenesis in symbiotic Aiptasia, that was not the case with aposymbiotic Aiptasia, where starvation led to increased NF-κB protein and increased susceptibility to bacteria-induced mortality. Therefore, NF-κB is clearly not the sole molecular inducer of immunity in Aiptasia. That is, NF-κB may have other or additional biological roles Aiptasia. Indeed, previous work^28^ showed that heat- or menthol-induced loss of symbionts in Aiptasia leads to increased levels of NF-κB, suggesting that algal symbionts suppress NF-κB activity in order to facilitate the establishment of symbiosis. Furthermore, NF-κB and several putative NF-κB-responsive genes are rapidly (within 3 h), albeit transiently, induced by heat shock in Aiptasia^51^. Of note, we have found that many of the same heat shock-induced genes with multiple NF-κB sites in their proximal promoters are also induced by starvation in Aiptasia. However, it has not been shown that these genes are direct target genes of Ap-NF-κB, nor have the specific roles of NF-κB or its target genes in the regulation of symbiosis, the response to heat shock, or starvation-induced biological responses been determined. Still, the induction of NF-κB under starvation in Aiptasia does suggest that this putative symbiosis-regulatory pathway is prioritized under energy-limiting conditions.

Overall, there is credible evidence that NF-κB has different roles in Aiptasia and *N. vectensis*. Similar to Aiptasia, the facultative coral *O. arbuscula* also shows an upregulation of immune response pathways in starved conditions. Moreover, thermal challenges, which have been well-documented to induce symbiont cell loss^8, 23, 61, 62^, have been shown to upregulate NF-κB transcripts in not only symbiotic anemones^51, 63^, but also in multiple species of symbiotic coral^64–67^. Thus, it appears that NF-κB has different biological roles in cnidarians that normally exist with vs. without symbionts. Furthermore, the shared expression of NF-κB and NF-κB target genes following both heat shock and starvation strengthens the argument for a role of this transcription factor pathway in affecting symbiont cell density under conditions that are unfavorable to the cnidarian-algal symbiosis. The conservation of NF-κB pathway induction between thermal stress and starvation supports a role for this transcription factor in the regulation of symbiont cell density under nutrient-poor conditions^25^. Our research contributes to the diversification of NF-κB biology beyond immunity, as has been seen in several organisms, including the control of early embryogenesis in *Drosophila* ^68^ and the development of cnidocytes in *N. vectensis*^69^.

In contrast to the NF-κB pathway and pathogen responses, starvation leads to increased susceptibility to oxidative stress as well as downregulation of oxidative stress response genes in both symbiotic and aposymbiotic states. For example, four SOD homologs were downregulated under starvation in Aiptasia from both states and an additional SOD homolog was downregulated in aposymbiotic Aiptasia only. SODs are involved in quenching reactive oxygen species, which are often produced by electron transport enzymes and increase in concentration under stress. This quenching ameliorates cell damage^70^. Increased expression of SOD genes has been documented under various stress conditions in Aiptasia both in and out of symbiosis^34, 71^ as well as in multiple species of coral^72^. These results demonstrate that, unlike the NF-κB pathway, energy is directed away from the ROS response under starvation and that the response to oxidative stress is not an energetic priority for Aiptasia under nutrient-limiting conditions.

The analysis of shared GO terms across three cnidarian species under starvation showcased differences in expression of immune and oxidative stress terms according to the organism’s capacity to establish symbiosis. Although oxidative stress terms were consistently underrepresented following starvation across all examined cnidarian taxa, immune terms exhibited variation; they were overrepresented following starvation in Aiptasia and *O. arbuscula* (both facultatively symbiotic species), while they were underrepresented in starved nonsymbiotic *N. vectensis*. These results provide evidence that a cnidarian’s capacity for symbiosis leads to distinct energetic priorities and regulatory pathways within the broader “defense” response. These results further demonstrate the need for caution when applying broad categorization of gene expression pathways that were described in other, evolutionarily distinct species, which may have evolutionarily distinct immune and stress regulatory pathways^73–75^.

Overall, this study demonstrates that cnidarian responses to pathogen or stress challenges are dependent on multiple factors, including symbiotic state, symbiotic capacity, and nutrition levels, which should be considered when predicting how cnidarians will respond in the face of perturbation^21^. Additionally, this study characterizes different energetic priorities between oxidative stress (low priority), symbiont-regulatory immunity (high priority), and pathogen immunity (dependent on symbiotic status and/or heterotrophic capacity) within the cnidarian “defense” response. The observation that pathogen immunity can depend on both symbiotic status and nutritional state may explain the contrasting effects of bleaching on pathogen immunity by accounting for the capacity of the bleached animal to supplement its energetic needs with heterotrophy^30, 76–80^. Furthermore, our findings contribute to the growing literature exploring the diverse biological processes (*e.g.,* immunity, development, and symbiosis) impacted by NF-κB in basal metazoans^20, 28, 53, 81^. The implications of these results are three-fold: 1) NF-κB and NF-κB gene pathways are either independent from Aiptasia pathogen-linked immunity, or they are not sufficient to defend against pathogen-induced mortality in the absence of adequate energetic reserves (as in starved aposymbiotic Aiptasia); 2) the oxidative stress response is dependent upon the availability of accessible nutrients beyond simple sugars; and 3) the oxidative stress response and the immune response have distinct energetic priorities in the facultative cnidarian Aiptasia.

The insights from the present study broaden our understanding of the mechanisms by which nutrient availability affect the stability of the cnidarian-algal symbiosis under various, compounding stressors^13^. Previous research has revealed that increased concentrations of inorganic compounds from anthropogenic sources compared to organic, bioavailable nutrient forms hinder photosynthesis and can disrupt nutrient exchange between endosymbionts and cnidarian hosts^13, 82, 83^. Furthermore, symbiont loss in episodes of coral bleaching results in starvation when hosts cannot effectively counterbalance nutrient loss with heterotrophy^84^. Thus, it is important to understand the molecular and physiological responses of cnidarians under conditions of nutrient availability and how these responses vary by species, symbiotic states, and symbiotic capacities.

## Acknowledgements

This research was supported by National Science Foundation grant IOS-1937650 (to T.D.G. and S.W.D.). M,V.-I. was supported by an NSF Graduate Research Fellowship and NSF NRT DGE 1735087. P.J.A.C. was supported by a Warren McLeod Marine Fellowship. C.B. and N.A.D.. were supported by the Boston University Undergraduate Research Opportunities Program, and N.A.D. received support from New England Biolabs and Dr. Loren Wold. We thank Stephen Lory (Harvard University) for *Pseudomonas aeruginosa* strain Pa14 and Kim Ritchie (University of South Carolina Beaufort) for *Serratia marcescens*.

