## Supplemental Tables and Figures for "Nutrient deprivation differentially affects gene expression, immunity, and pathogen susceptibility across symbiotic states in a model cnidarian"

**This file includes:**

Supplementary Figures 1 and 2

Supplementary Tables 1, 2, 3

High resolution versions of all supplementary materials, GO term dendrograms, as well as scripts, *Exaiptasia pallida* reference files, and all data files and figures generated for this manuscript are publicly available at: <https://github.com/mariaingersoll/Aiptasia_Fed_Starved.git>

**Other materials for this manuscript include the following:**

Supplementary Dataset 1

Supplementary Dataset 2

Supplementary Dataset 3

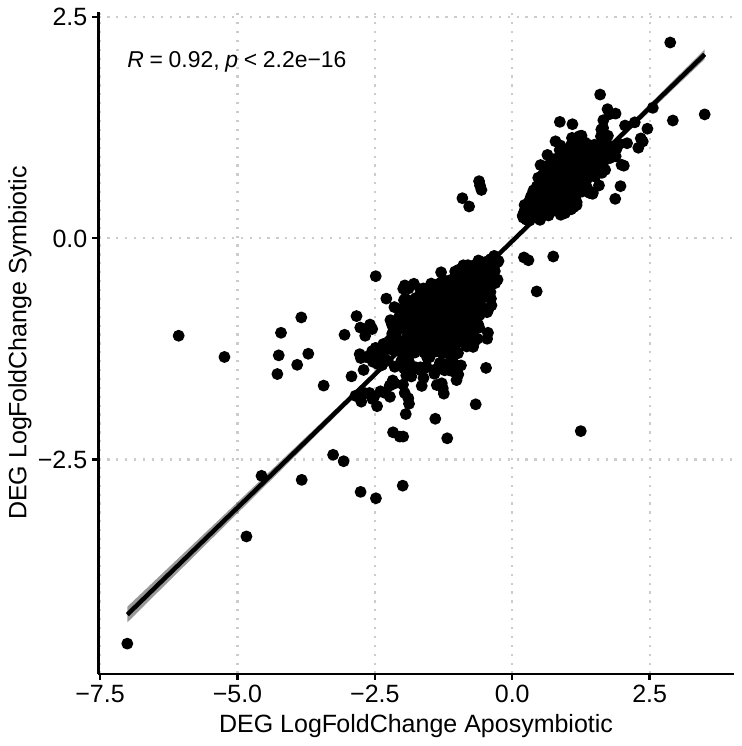

**Supplementary Figure 1.** Comparison of log2 fold changes (LogFoldChange) reveals a positive correlation (R=0.92, p<2.2e-16) in the expression of all 1,466 shared Differentially Expressed Genes (DEGs; FDR p-value <0.05, **Fig. 2C**) following starvation of aposymbiotic (x axis) and symbiotic (y axis) Aiptasia.

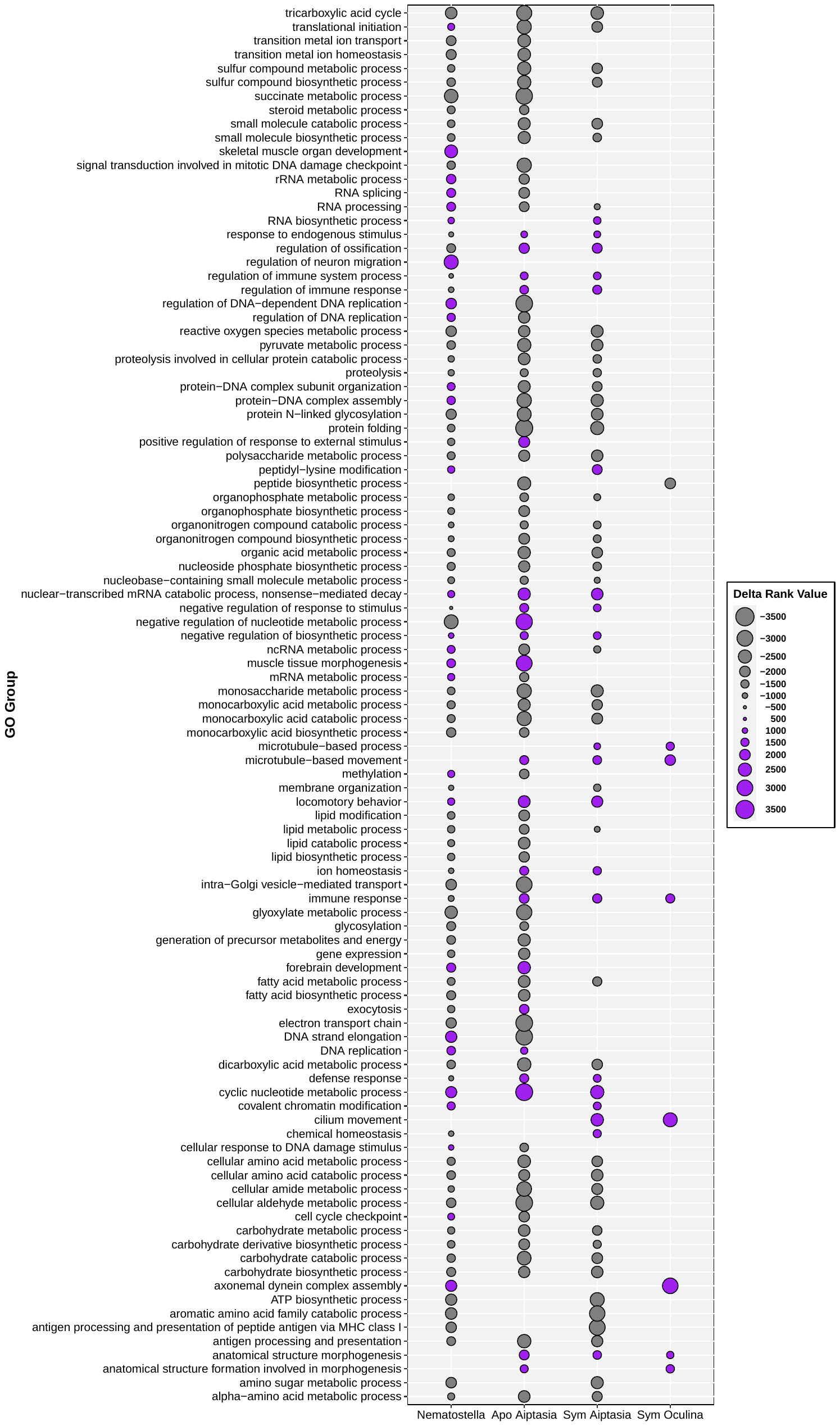

**Supplementary Figure 2.** Delta rank values (represented by point sizes) of all GO terms significantly enriched in fed (grey, negative delta rank values) or in starved (purple, positive delta rank values) treatments of at least two cnidarians (*Nematostella vectensis* (Aguirre Carrión et al., 2023), aposymbiotic Aiptasia, symbiotic Aiptasia, or symbiotic *Oculina arbuscula* (Rivera et al., 2023)). GO terms enriched in aposymbiotic and symbiotic Aiptasia, but not in either of the other species, were not included.

**Supplementary Table 1.** Differentially Expressed Genes (DEGs) regulated in opposite directions in symbiotic and aposymbiotic Aiptasia following starvation. Positive log2 fold change (LFC) values indicate higher expression following starvation compared to fed controls. All LFC values have been rounded to two decimal places.

| **Gene** | **Gene Symbol** | **Annotation** | **ApoFS**  **LFC** | **SymFS**  **LFC** |
| --- | --- | --- | --- | --- |
| *Downregulated in aposymbiotic, upregulated in symbiotic* | | | | |
| AIPGENE  12535 | CTHRC1.4 | Collagen triple helix repeat-containing protein 1 | -0.59 | 0.59 |
| AIPGENE  13828 | LACS8 | Long chain acyl-CoA synthetase 8 | -0.91 | 0.45 |
| AIPGENE  17478 | ASNS | Asparagine synthetase [glutamine-hydrolyzing] | -0.79 | 0.36 |
| AIPGENE  2827 | HYPOTHETICAL.31 | Hypothetical protein NEMVEDRAFT_v1g248480 | -0.60 | 0.64 |
| AIPGENE  4127 | CARBOXYPEPTIDASE | Carboxypeptidase inhibitor SmCl | -0.56 | 0.54 |
| *Upregulated in aposymbiotic, downregulated in symbiotic* | | | | |
| AIPGENE  21459 | CAT.1 | Catalase | 0.75 | -0.21 |
| AIPGENE  21713 | THYMOSIN | Thymosin beta-b | 0.21 | -0.22 |
| AIPGENE  22059 | CYB5A.1 | Cytochrome b5 | 1.25 | -2.18 |
| AIPGENE  23774 | TVAG_198570.1 | Viral A-type inclusion protein, putative | 0.29 | -0.25 |
| AIPGENE  4821 | BBOX1 | Gamma-butyrobetaine dioxygenase | 0.44 | -0.60 |

**Supplementary Table 2:** Heat and starvation responsive putative NF-κB target genes from Figure 4B. Gene names, symbols, and annotation from the *Exaiptasia pallida* (Aiptasia; Baumgarten et al., 2015), as well as the number of putative NF-κB binding sites in the gene promoter region (500 bp upstream of the transcription start site) from Cleves et al. (2020). Genes with no annotation are marked with n.a.

| **Gene** | **Gene Symbol** | **Annotation** | **NF-κB Sites** |
| --- | --- | --- | --- |
| AIPGENE  18569 | IFIH1.2 | Interferon-induced helicase C domain-containing protein 1 | 3 |
| AIPGENE  18872 | BCL3 | B-cell lymphoma 3 protein | 6 |
| AIPGENE  18937 | MAFG | Transcription factor MafG | 6 |
| AIPGENE  23318 | GBP5.1 | Guanylate-binding protein 5 | 5 |
| AIPGENE  26161 | MAF | Transcription factor Maf | 3 |
| AIPGENE  10002 | ELF3 | ETS-related transcription factor Elf-3 | 3 |
| AIPGENE  10155 | IFIH1 | Interferon-induced helicase C domain-containing protein 1 | 4 |
| AIPGENE  12036 | X.1657 | n.a. | 6 |
| AIPGENE  12505 | X.1741 | n.a. | 3 |
| AIPGENE  17157 | ACD_16C00100G0095 | Tetratricopeptide repeat protein | 3 |
| AIPGENE  27025 | X.3853 | n.a. | 5 |
| AIPGENE  27057 | RCHY1.1 | RING finger and CHY zinc finger domain-containing protein 1 | 7 |
| AIPGENE  8848 | NFKB1 | Nuclear factor NF-kappa-B p105 subunit | 3 |

**Supplementary Table 3:** Expression levels of heat and starvation responsive putative NF-κB target genes from Figure 4B. Log2 fold change (LFC) and FDR p-value (padj) of each gene following starvation in aposymbiotic (ApoFS) and symbiotic (SymFS) Aiptasia, as well as LFC and padj of each gene following three hours of heat shock in aposymbiotic (ApoHeat) and symbiotic (SymHeat) Aiptasia (from Cleves et al., 2020). Positive LFC values indicate higher expression following treatment (starvation or heat shock) compared to control conditions (feeding or ambient temperature). All numerical values have been rounded to two decimal places. Genes that were not significantly expressed following starvation in symbiotic Aiptasia are noted with n.s. in SymFS LFC and SymFS padj columns.

| **Gene** | **ApoFS**  **LFC** | **ApoFS**  **padj** | **SymFS**  **LFC** | **SymFS**  **padj** | **ApoHeat**  **LFC** | **ApoHeat**  **padj** | **SymHeat**  **LFC** | **SymHeat padj** |
| --- | --- | --- | --- | --- | --- | --- | --- | --- |
| *Heat responsive putative NF-κB target genes induced by starvation in apo and sym Aiptasia* | | | | | | | | |
| AIPGENE  18569 | 0.91 | 7.87E-05 | 0.61 | 1.31E-02 | 4.39 | 0 | 3.85 | 0 |
| AIPGENE  18872 | 1.17 | 6.12E-22 | 0.41 | 3.72E-03 | 2.50 | 0 | 2.02 | 0 |
| AIPGENE  18937 | 1.25 | 1.91E-12 | 0.69 | 3.91E-04 | 3.09 | 0 | 2.50 | 0 |
| AIPGENE  23318 | 1.78 | 1.09E-04 | 1.40 | 4.48E-03 | 3.76 | 0 | 3.04 | 0 |
| AIPGENE  26161 | 0.58 | 1.11E-02 | 0.63 | 6.77E-03 | 4.64 | 0 | 3.89 | 0 |
| *Heat responsive putative NF-κB target genes induced by starvation in only apo Aiptasia* | | | | | | | | |
| AIPGENE  10002 | 0.38 | 2.13E-02 | n.s. | n.s. | 3.08 | 0 | 2.22 | 0 |
| AIPGENE  10155 | 1.31 | 2.30E-02 | n.s. | n.s. | 1.87 | 0 | 2.56 | 0 |
| AIPGENE  12036 | 0.61 | 2.14E-02 | n.s. | n.s. | 2.60 | 0 | 2.61 | 0 |
| AIPGENE  12505 | 0.61 | 2.57E-04 | n.s. | n.s. | 2.85 | 0 | 2.13 | 0 |
| AIPGENE  17157 | 1.10 | 2.57E-03 | n.s. | n.s. | 3.15 | 0 | 2.81 | 0 |
| AIPGENE  27025 | 0.55 | 7.74E-03 | n.s. | n.s. | 3.24 | 0 | 3.08 | 0 |
| AIPGENE  27057 | 0.71 | 1.05E-02 | n.s. | n.s. | 3.93 | 0 | 3.22 | 0 |
| AIPGENE  8848 | 1.21 | 3.96E-03 | n.s. | n.s. | 4.20 | 0 | 2.46 | 0 |
